# Galectin-3 Mediated Endocytosis of the Orphan G-Protein-Coupled Receptor GPRC5A

**DOI:** 10.1101/2025.08.06.668629

**Authors:** Abdeldjalil Boucheham, Jorge Mallor Franco, Séverine Bär, Ewan MacDonald, Solène Zuttion, Lana Blagec, Bruno Rinaldi, Johana Chicher, Laurianne Kuhn, Philippe Hamman, Christian Wunder, Ludger Johannes, Hocine Rechreche, Sylvie Friant

## Abstract

Galectins, a family of glycan-binding proteins, play crucial roles in various cellular functions, acting at both intracellular and extracellular levels. Among them, Galectin-3 (Gal-3) stands out as a unique member, possessing an intrinsically disordered N-terminal interaction domain and a canonical carbohydrate-recognition domain (CRD). Gal-3 binding to glycosylated plasma membrane cargo leads to its oligomerization and membrane bending, ultimately resulting in the formation of endocytic invaginations. An interactomic assay using proteomic analysis of endogenous Gal-3 immunoprecipitates identified the orphan G protein-coupled receptor GPRC5A as a novel binding partner of Gal-3. GPRC5A, also known as Retinoic Acid-Induced protein 3 (RAI3), is transcriptionally induced by retinoic acid. Our results further demonstrate that extracellular recombinant Gal-3 stimulates GPRC5A internalization. In SW480 colorectal cancer cells, glycosylated GPRC5A interacts with Gal-3. Interestingly, while GPRC5A expression was upregulated by the addition of all-trans retinoic acid (ATRA), its endogenous internalization in SW480 cells was specifically triggered by extracellular Gal-3, but not by ATRA. This study provides new insights into the endocytic mechanisms of GPRC5A, for which no specific ligand has been identified to date. Further research may uncover additional Gal-3 mediated functions in GPRC5A cellular signaling and contribute to the development of innovative therapeutic strategies.

## 1. Introduction

Galectins are glycan-binding proteins (lectins) that bear at least one carbohydrate-recognition domain (CRD), which binds to β-galactosides. Galectins can function intracellularly and can also be secreted to bind to cell surface glycoconjugated receptors. Galectins are involved in many cellular functions, such as cell-matrix adhesion, cell interactions, signaling and membrane trafficking [1]. Given their dual extracellular and intracellular functions, galectins are regarded as major players in normal physiology as well as in different disease conditions such as oncogenic processes. Structurally, there are three subfamilies of mammalian galectins: prototype, tandem repeat and chimera. Galectin-3 (Gal-3) encoded by the *LGALS3* gene is the only member of the chimera group. Gal-3 is a 26 kDa lectin with a disordered N-terminal domain involved in inter- and intra-molecular interactions and a single CRD [2]. Expression of *LGALS3* gene is altered in many types of cancer, and Gal-3 is identified as an effective drug target for cancer diagnostics, as well as for inflammatory and fibrotic diseases [3-5]. Recent data show that increased blood concentrations of Gal-3 are observed in patients with severe COVID-19, highlighting its importance as a prognostic factor and as a promising treatment target [6,7]. Gal-3 multimerization via the N-terminal domain results in clustering, a process promoted by Gal-3 binding to glycan ligands through its CRD [1]. Moreover, Gal-3 binding to a specific glycoproteins and glycolipids allows rearrangement of their organization in the membrane [8]. Gal-3 critical role in endocytosis is achieved via its recruitment to membranes by binding glycosylated cargos such as CD44, CD98, β1-integrin, or lactotransferrin, next clustering of these cargo proteins and glycosphingolipids generates membrane bending and the formation of endocytic invaginations [9-12]. This galectin-dependent endocytic mechanism has been termed the GlycoLipid-Lectin (GL-Lect) endocytosis [13,14].

G protein-coupled receptors (GPCRs) constitute a large family of conserved membrane receptors with seven transmembrane domains. They are activated by a diverse range of extracellular molecules to mediate signal transduction pathways. Furthermore, their involvement in many physiological and pathophysiological cellular processes establishes them as the most common drug targets [15]. Based on sequence homology, several receptor subfamilies have been identified. Among them, the GPCR family C group 5 (GPRC5) receptors are classified as a subfamily with four members, GPRC5A-D, for which no specific ligands are described. GPRC5A was identified as an all-*trans*-retinoic acid (ATRA) up-regulated gene initially termed RAIG1 retinoic-acid inducible gene 1, and later also termed RAI3 [16]. A recent chemoproteomic profiling study shows that microbiota-derived aromatic monoamines can bind to GPRC5A and stimulate GPRC5A–β-arrestin recruitment [17]. Increased expression of GPRC5A is associated with colon, pancreas, and prostate cancers and could serve as a candidate biomarker for accurate diagnosis and prognosis [18,19].

Building on the established role of Gal-3 in mediating endocytosis of membrane-associated proteins, we performed a Gal-3 interaction study using mass spectrometry to identify potential Gal-3 binding partners. The identified GPRC5A was internalized after the addition of Gal-3 and might follow a similar mechanism to those reported for other Gal-3 binding partners.

## 2. Materials and Methods

### 2.1. Cell culture

The HepG2 (hepatocarcinoma) and HeLa (cervical adenocarcinoma) cell lines, along with the colorectal adenocarcinoma cell lines SW480, Caco-2, DLD1, HT29, and HCT116, were kindly provided by Dr. Isabelle Gross (CRBS, Strasbourg). Cells were cultured in DMEM supplemented with 10% fetal calf serum (FCS), 1% penicillin-streptomycin, and L-glutamine (Gibco). Cultures were maintained at 37°C in a humidified incubator with 5% CO2 and were passaged twice a week. For immunofluorescence, cells were seeded one day before the experiments. Transient transfection was performed using Lipofectamine® 2000 (ThermoFisher) according to the manufacturer’s instructions.

### 2.2. Purification of recombinant Galectin-3

Human recombinant wild-type Gal-3 with C-terminal 6xHis tags was prepared as previously described [12]. Briefly, Gal-3 expression was induced at 20°C in Rosetta2-pLysS using three L of LB-media with 60 μM IPTG overnight, and purified by cobalt resin (Thermo Fisher Scientific) affinity chromatography and gel filtration (Superdex75 10×30) in PBS at pH 7.3. Small aliquots for single use were snap-frozen and stored at -80 °C.

### 2.3. Immunofluorescence

Immunofluorescence assays were performed as previously described [20]. For immunofluorescence, cells were grown on coverslips in Nunc 4-well or 6-well culture plates (ThermoFisher). Cells were fixed with 4% paraformaldehyde for 10 minutes at room temperature and permeabilized with 0,2% Triton-X100 for 10 minutes. Following three washes with 1× PBS, cells were blocked with 20% FCS in 1× PBS for 1 hour at room temperature (or overnight at 4°C). After three additional washes, primary antibodies directed against GPRC5A (MilliporeSigma, HPA046526) were diluted in 2% FCS-1X PBS and applied to the samples, followed by incubation with IgG-conjugated AlexaFluor 568 secondary antibodies (ThermoFisher) for 1h. After a final washing step, coverslips were mounted with Vectashield No-Fade mounting medium containing DAPI (Vector laboratories). Imaging was performed using a Zeiss Axio Observer D1 fluorescence microscope or Zeiss LSM700 confocal microscope (40× objective, Plateforme Microscopie et Imagerie, IBMP, Strasbourg, France or Plateforme d’Imagerie du CRBS PIC-STRA, Strasbourg, France), and images were analyzed using ImageJ.

### 2.4. Cell lysis for total protein extraction

Confluent cells were washed twice with ice-cold 1× PBS and lysed on ice for 5 mins in a buffer containing 50 mM Tris-HCl (pH 8), 50 mM NaCl, 1% NP-40, and a cOmplete EDTA-free protease inhibitor cocktail (MilliporeSigma). Cells were scrapped and incubated for 5 mins on ice; the resulting lysates were centrifuged at 14000 g for 15 mins at 4°C. The collected supernatant (total protein extract) was quantified using Bradford Protein Assay reagent (BioRad).

### 2.5. Immunoprecipitation experiments and western blot analyses

Two different immunoprecipitation methods were used to pull down proteins: µMACs Protein G magnetic beads isolation kit (Miltenyi Biotec) and Gamma-Bind Plus Sepharose beads (Cytiva). The µMACS protocol was chosen for its magnetic separation efficiency, while the Gamma-Bind protocol provided an alternative for confirmation of protein interactions. Cells were washed with ice-cold 1× PBS, scraped into tubes, and lysed on ice for 5 mins with lysis buffer (50 mM Tris-HCl, 50 mM NaCl, 1% NP-40, supplemented with cOmplete EDTA-free protease inhibitor cocktail, MilliporeSigma). The lysate was clarified by centrifugation at 13, 000 g for 13 mins at 4°C. For the µMACS protocol, 1.2 mg of protein lysate was incubated with 50 µl of µMACS Protein G magnetic microbeads (Miltenyi Biotec) coupled to anti-Gal-3 antibodies (Mouse monoclonal, Santa Cruz sc-32790). After 1 hour of incubation at 4°C, samples were loaded onto a µMACS Column in the magnetic field of a µMACS Separator. Beads and associated proteins were retained during extensive washing steps (3 × 500 µl lysis buffer washes), and elution was performed with 100 µl of elution buffer provided in the kit. Eluded proteins were identified by mass spectrometry.

For Gamma-Bind assays, Gamma-Bind Plus Sepharose beads were equilibrated with lysis buffer and incubated with anti-Gal-3 antibodies (Mouse monoclonal, Santa Cruz sc-32790) at 4°C for 2 hours. After washing (3 × 500 µl lysis buffer) 1.5 mg of protein lysate was added to the beads and incubated at 4°C for 1 hour with gentle agitation. After extensive washing steps, elution was performed with 5× Laemmeli buffer, followed by incubation at 37°C for 5 mins. After centrifugation, the resulted supernatant was analyzed by SDS-PAGE. 2,2,2-Trichloroethanol (TCE) incorporated into SDS-PAGE gels enabled stain-free fluorescent detection of proteins [21]. Proteins were resolved using SDS–PAGE and transferred to a Protran® nitrocellulose membrane (Amersham). Membranes were blocked with 4% non-fat milk in PBST for 1 hour at room temperature, then incubated with primary antibodies: anti-GPRC5A (1:1,000, MilliporeSigma, HPA007928), anti-Gal-3 (1:2,000, Abcam, ab76245), anti-RIP (1:1,000, BD Biosciences, AB_397832), and anti-GAPDH (1:10,000; Abcam rabbit #ab181602) overnight at 4°C. After washing (3 × 10 minutes in PBST), membranes were incubated with HRP-conjugated secondary antibodies (1:10,000; ThermoFisher) for 1 hour at room temperature. Proteins were detected using ECL substrate (ThermoFisher), and images were acquired with a ChemiDocTouch imaging system (BioRad).

### 2.6. Mass spectrometry analysis

Protein extracts were prepared as described in a previous study [22]. Each sample was precipitated with 0.1 M ammonium acetate in 100% methanol, and proteins were resuspended in 50 mM ammonium bicarbonate. After a reduction-alkylation step (dithiothreitol 5 mM and iodoacetamide 10 mM), proteins were digested overnight with sequencing-grade porcine trypsin (1:25, w/w, Promega). The resulting vacuum-dried peptides were resuspended in water containing 0.1% (v/v) formic acid (solvent A). One sixth of the peptide mixture were analyzed by nanoLC-MS/MS an Easy-nanoLC-1000 system coupled to a Q-Exactive Plus mass spectrometer (ThermoFisher) operating in positive mode. Five microliters of each sample were loaded on a C-18 precolumn (75 μm ID × 20 mm nanoViper, 3µm Acclaim PepMap; ThermoFisher) coupled with the analytical C18 column (75 μm ID × 25 cm nanoViper, 3µm Acclaim PepMap; ThermoFisher). Peptides were eluted with a 160 min gradient of 0.1% formic acid in acetonitrile at 300 nL/min. The Q-Exactive Plus was operated in data-dependent acquisition (DDA) mode with Xcalibur software (ThermoFisher). Survey MS scans were acquired at a resolution of 70.000 at 200 m/z (mass range 350-1250), with a maximum injection time of 20 ms and an automatic gain control (AGC) set to 3e6. Up to 10 of the most intense multiply charged ions (≥2) were selected for fragmentation with a maximum injection time of 100 ms, an AGC set at 1e5 and a resolution of 17.500. A dynamic exclusion time of 20 s was applied during the peak selection process.

### 2.7. Database search and mass spectrometry data post-processing

MS data were searched against the UniProtKB database (release 2016_08, 149870 forward sequences) with Human taxonomy. We used the Mascot algorithm (version 2.3, Matrix Science) to perform the database search with a decoy strategy and search parameters as follows: carbamidomethylation of cysteine was set as fixed modification; N-terminal protein acetylation, phosphorylation of serine / threonine / tyrosine and oxidation of methionine were set as variable modifications; tryptic specificity with up to three missed cleavages was used. The mass tolerances in MS and MS/MS were set to 10 ppm and 0.02 Da, respectively, and the instrument configuration was specified as “ESI-Trap”.

The resulting .dat Mascot files were then imported into Proline v1.4 package (http://proline.profiproteomics.fr/) for post-processing. Proteins were validated with Mascot pretty rank equal to 1, and a 1% false discovery rate (FDR) on both peptide spectrum matches (PSMs) and protein sets (based on score). The total number of MS/MS fragmentation spectra was used to quantify each protein in the different samples. The co-immunoprecipitation data were compared with the data collected from multiple experiments against the negative control to identify significant differences, as previously described [23].

Mass spectrometry data analysis and visualization were performed using R version 4.4.3 [24] within the RStudio integrated development environment (version 2024.12.1.563; [25]). The following packages: tidyverse [26], ggplot2 [27], readxl [28], openxlsx [29], and ggrepel [30] were used for data manipulation and plotting, Excel import/export, and non-overlapping label annotation. Proteins were quantified based on the number of identified spectra per replicate and for each protein; the mean peptide count was calculated across biological replicates in both anti-Gal-3 IP experiment (n=4) and negative control without antibodies (n=4) conditions. To compute the log_²_ fold change (log_²_FC), a pseudo-count of 0.1 was added to all values prior to ratio calculation in order to avoid division by zero or log transformation of zero values. An unpaired two-tailed Student’s t-test was performed for each protein to assess statistical significance between control and test conditions. The resulting p-values were transformed to –log_10_(p-value) for visualization, and proteins were considered significantly enriched if they met both criteria: log_²_FC ≥ 1 and p-value < 0.05. Finally, volcano plots were generated to visualize fold change against statistical significance. Thresholds for significance (log_²_FC ≥ 1 and p < 0.05) were marked with dashed blue (vertical) and green (horizontal) lines, and significantly enriched proteins were highlighted in red (Figure 1A).

**Figure 1.**
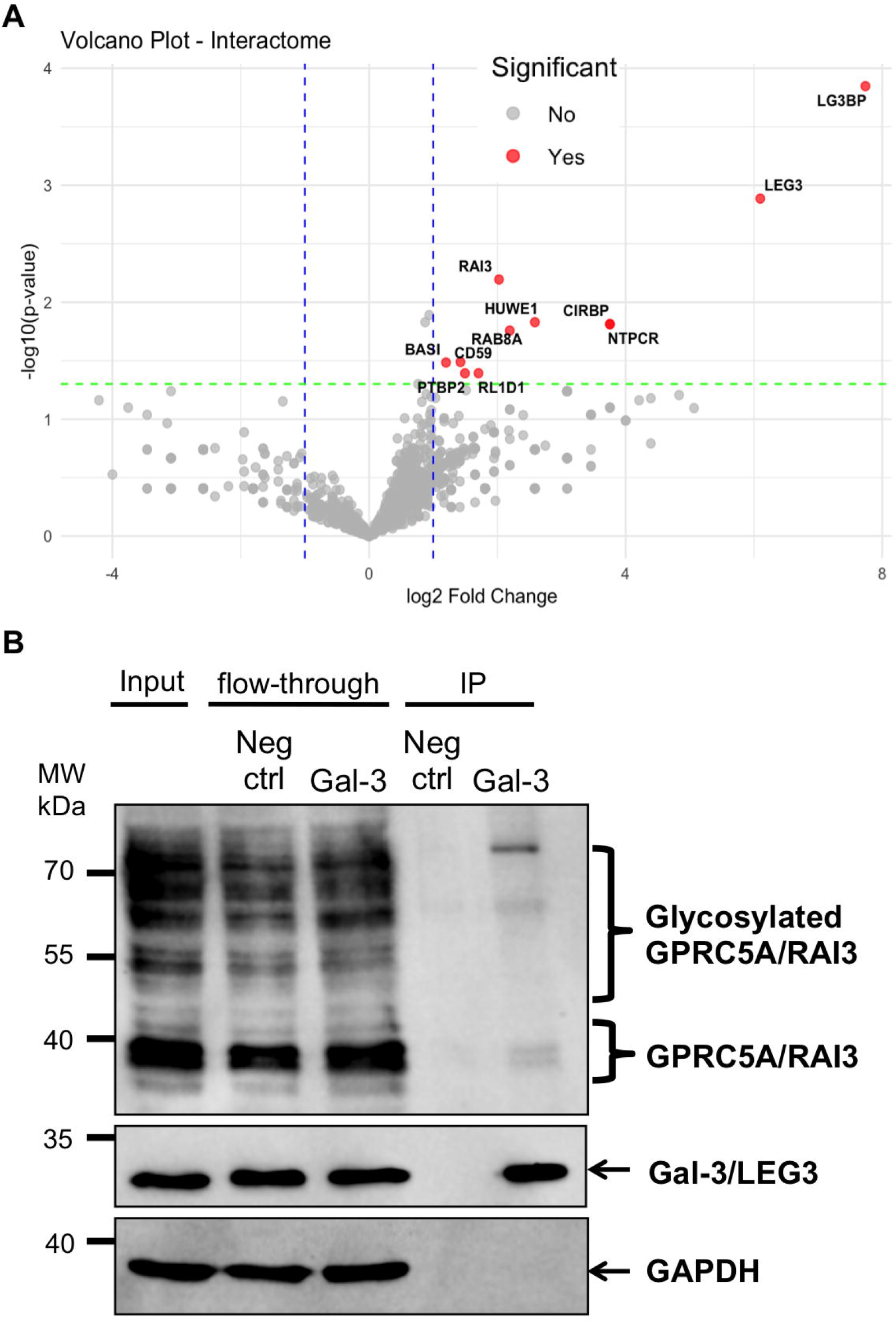
Identification of GPRC5A as a Gal-3 partner in HepG2 cells (**A**) Interactomic analysis of Gal-3 in HepG2 cells. Immunoprecipitation (IP) using anti-Gal-3 antibodies or a negative control (no antibody) was followed by nanoLC-MS/MS analyses. The results show the total number of identified spectra per protein across the different experiments. A volcano plot of –log_10_(p-value) versus log_²_(fold change) was generated to highlight proteins exhibiting significant differential detection between the IP (n = 4) and control (n = 4) conditions. Thresholds for significance (log_²_FC ≥ 1 and p < 0.05) are marked with dashed blue (vertical) and green (horizontal) lines, and significantly enriched proteins are highlighted in red. (**B**) Co-immunoprecipitation (co-IP) of Gal-3 (also termed LEG3) and GPRC5A (also termed RAI3). Total protein lysates (Input) were subjected to IP with anti-Gal-3 antibodies using µMACS Protein G magnetic microbead. The negative control (no antibody) was done in parallel. Immunoprecipitated (IP) and flow-through fractions were analyzed via Western blot with the indicated antibodies. The glycosylated forms of GPRC5A are indicated. GAPDH was used as a negative control.

The mass spectrometry proteomics data have been deposited to the ProteomeXchange Consortium via the PRIDE partner repository [31], under the dataset identifier PXD048507 and 10.6019/PXD048507.

### 2.8. Gal-3 dependent endocytosis assays

Non-confluent SW480 cells were incubated in cold DMEM containing 0,2% BSA and anti-GPRC5A antibody (MilliporeSigma, HPA046526) added for 10 minutes. Unbound antibody was removed by washing with PBS++ (1× PBS, 0.5 mM CaCl2, 0.5 mM MgCl2), followed by a rinse with ice-cold DMEM containing 2% BSA. To initiate endocytosis, cells were treated with 1 µg/mL of purified recombinant Gal-3 in pre-warmed DMEM supplemented with 0,2% BSA and incubated at 37°C for the indicated times. At each time point, the medium was replaced with PBS++, and cells were incubated on ice. To remove surface-bound antibodies or Gal-3, subsequent washes were performed three times with 0.5 M glycine (pH 2.2), followed by a single wash with 200 mM lactose. Cells were prepared for imaging using the Immunofluorescence protocol for GPRC5A (see section 2.3). The methodology is adapted from previously published works [9,20].

## 3. Results

### 3.1. Identification of GPRC5A as a Gal-3 binding partner

To identify binding partners of Gal-3, we performed an interactomics analysis of endogenous Gal-3 using an approach previously applied to the VPS15 membrane trafficking and autophagy effector [32]. HepG2 cells were lysed and subjected to immunoprecipitation (IP) with anti-Gal-3 antibodies or without antibodies as a negative control (n=4). After extensive washing, the bound proteins were identified using mass spectrometry. The interactomics data were validated by spectral counts and statistical analysis (Supplementary Figure 1). A scatterplot of -log10(p-value) against log2(fold change) was generated (Figure 1A) to visualize differentially enriched proteins. This analysis revealed significant enrichment of Galectin-3 (Gal-3, also known as LEG3) in the anti-Gal-3 IP samples compared to negative controls. Notably, this analysis also showed a significant enrichment of previously identified Gal-3 interacting partners, including Galectin-3 Binding Protein (Gal-3BP) [33], CD59 [34], and BASI (Basigin/CD147) [35]. Gal-3BP, also known as LGALS3BP, is a tumor-associated antigen of approximately 90 kDa. It is an extracellularly secreted glycoprotein and a well-characterized ligand of Gal-3 [33]. CD59, a GPI-anchored protein also known as Protectin, is an immunoregulatory receptor that protects human cells from complement-mediated damage [36]. BASI (Basigin), also known as CD147 or extracellular matrix metalloproteinase inducer (EMMPRIN), is a transmembrane glycoprotein belonging to the immunoglobulin superfamily and is highly expressed in human tumors [37]. These previously reported interactors associate physically with Gal-3 in a carbohydrate-dependent manner and contribute to immune modulation (Gal-3BP) [38], complement resistance (CD59) [34], and matrix remodeling (BASI/CD147) [39]. For the other significantly enriched interacting proteins, no experimental validation has been reported to date.

Focusing on transmembrane proteins, since Gal-3 is known to specifically interact with membrane receptors [13], our interactomic data also identified the orphan G-protein coupled receptor GPRC5A, known as RAI3. This receptor, whose expression is induced by retinoic acid, was significantly enriched in the Gal-3 IP samples. To confirm the interaction, we performed a co-IP assay (Figure 1B), which revealed that Gal-3 (26 kDa) interacts with both the 40 kDa form of GPRC5A as well as a higher molecular weight form, previously described as glycosylated [40]. Although the interaction was confirmed, the overall abundance of GPRC5A in the IP fractions was low. Notably, however, a higher molecular weight form of GPRC5A, likely corresponding to its glycosylated form, was more prominent in the IP fraction (Figure 1B). Consequently, we extended our analysis to additional cell types to investigate the interaction under conditions showing higher levels of GPRC5A glycosylated forms. This strategy is consistent with findings by Greenhough and colleagues [41], who reported that GPRC5A exhibits multiple electrophoretic migration bands.

### 3.2. Increased GPRC5A level and interaction with Gal-3 in colon cancer cells

The expression level of the *GPRC5A* gene varies across cancer types, displaying increased expression in colon, pancreas, and prostate cancers, and decreased expression in lung cancer [19]. Accordingly, we analyzed GPRC5A and Gal-3 protein levels compared to the RIP protein (loading control) and a TCE staining (loading control) in cell lines derived from colorectal (SW480, HCT116, HT-29, DLD-1, Caco-2), gastric (AGS) and cervical adenocarcinoma (HeLa) cancers, as well as HepG2 hepatocarcinoma cells (Figure 2A). Compared with HepG2 cells, most of these cancer cell lines, except the gastric AGS cells, exhibited higher levels of GPRC5A, especially the higher molecular weight forms previously reported to be glycosylated [40]. From these, we selected SW480 human colon adenocarcinoma cells and tested the interaction between Gal-3 and GPRC5A by co-IP (Figure 2B). We observed interaction of Gal-3 with GPRC5A, with a stronger signal for the high-molecular weight forms compared to the 40 kDa form. These results indicate that in cancer cells, the carbohydrate-binding lectin Gal-3 binds to glycosylated GPRC5A.

**Figure 2.**
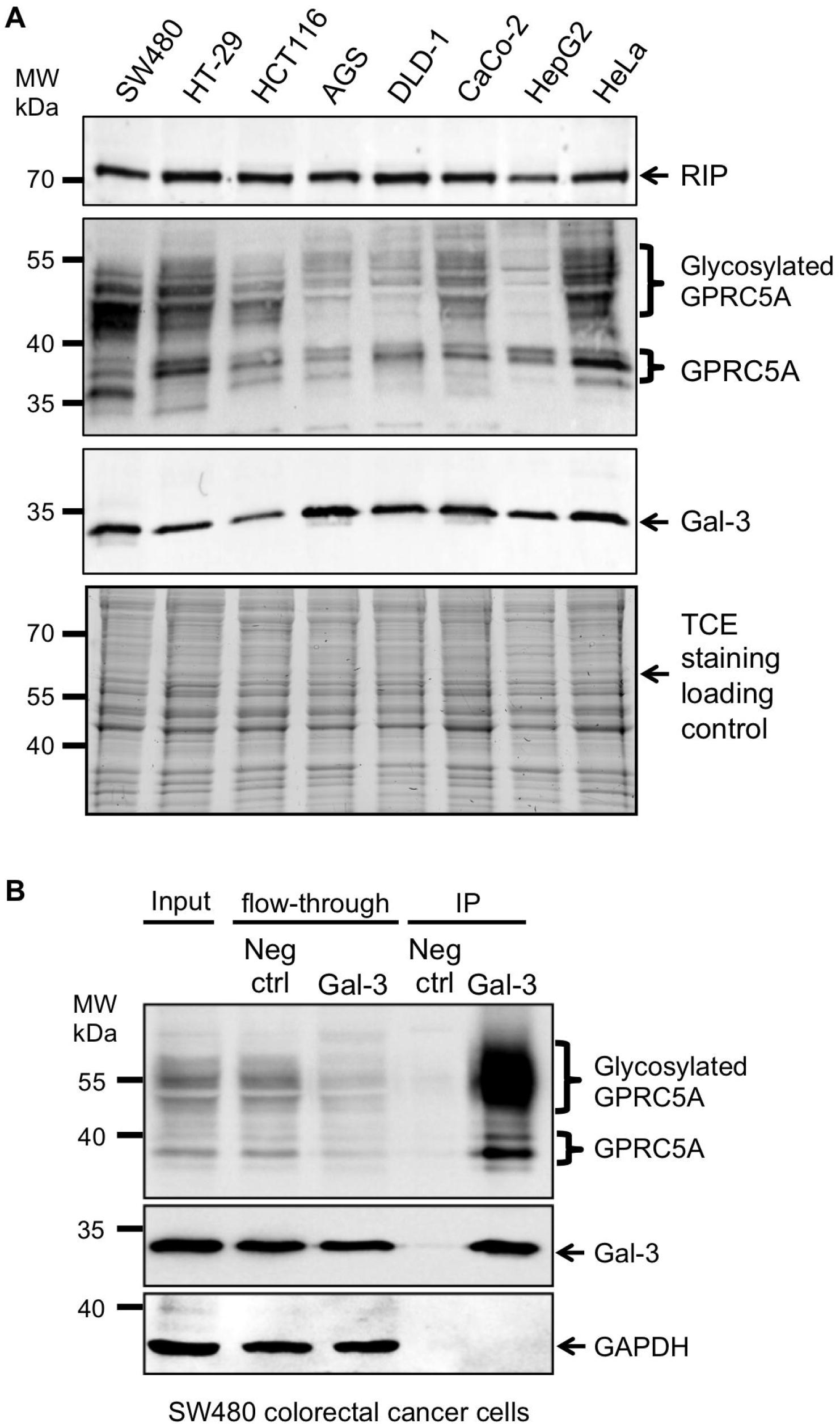
(A) GPRC5A and Gal-3 expression levels across cell lines. Western blot analysis using GPRC5A and Gal-3 antibodies and anti-RIP and TCE staining as loading controls were performed on total protein extracts from SW480, HT-29, HCT116, AGS, DLD-1, CaCo-2, HepG2, and HeLa cell lines. SW480 cells displayed prominently glycosylated GPRC5A forms. (B) Interaction between Gal-3 and GPRC5A in SW480 cells. Anti-Gal-3 immunoprecipitation (IP) or negative control IP without antibodies (Neg ctrl) was performed on total protein extracts (Input) from SW480 cells. Western blot analysis was performed on fractions (Input, Flow-through, and IP) using antibodies against GPRC5A, Gal-3, or GAPDH (negative control).

### 3.3. Addition of retinoic acid leads to increased levels of GPRC5A proteins

GPRC5A was previously characterized as an all-trans-retinoic acid (ATRA)-upregulated gene [16]. To assess the effect of ATRA on GPRC5A protein expression, we treated HepG2 (Figure 3A) and SW480 (Figure 3B) cells with ATRA (10^−6^ M-10^−6^ M) and analyzed GPRC5A and Gal-3 protein levels and profiles. Treatment with ATRA increased GPRC5A expression, in both cell types. Additionally, glycosylated GPRC5A forms, which strongly interact with Gal-3, were further upregulated in SW480 cells.

**Figure 3.**
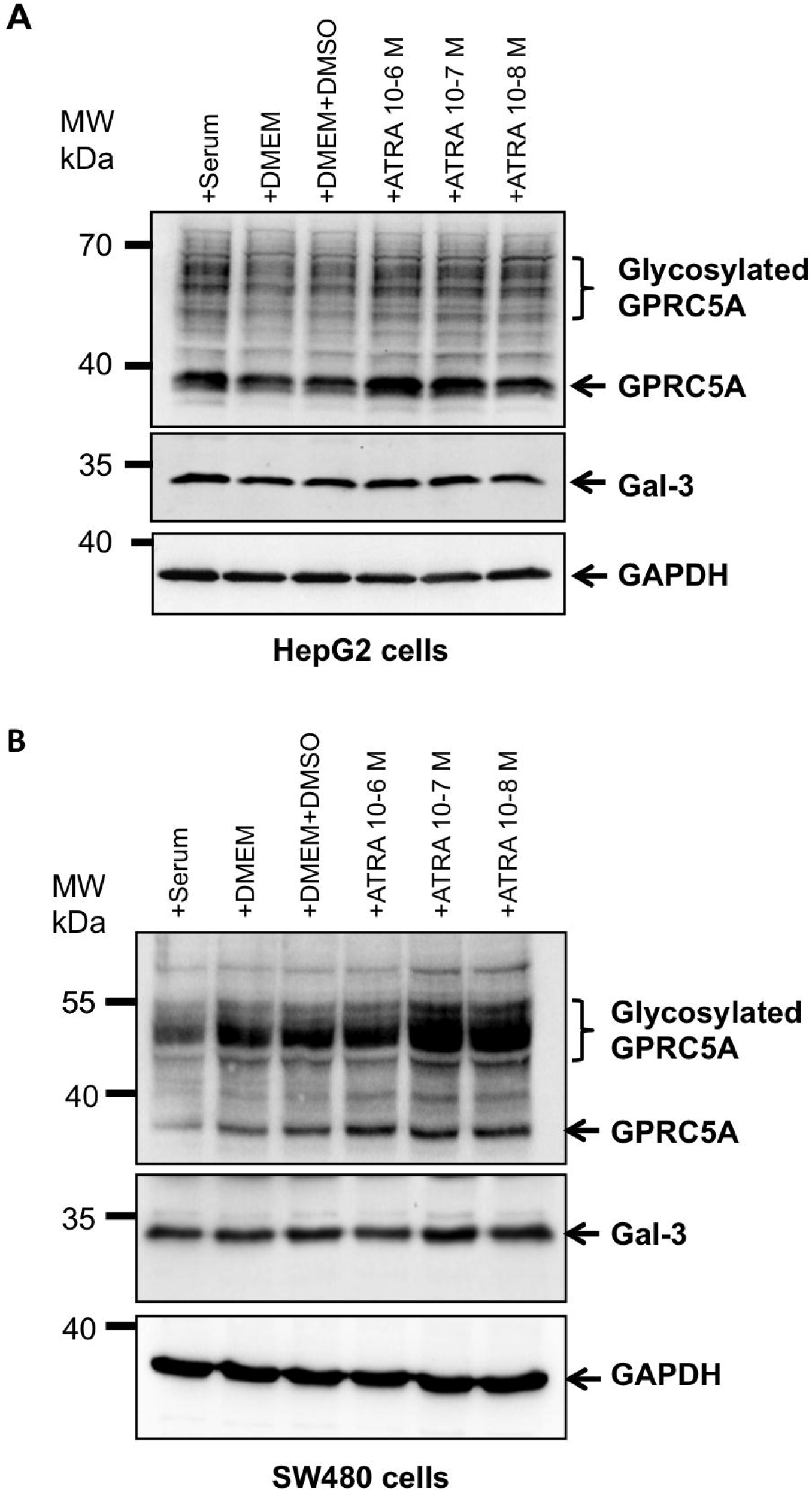
GPRC5A and Gal-3 protein levels upon retinoic acid addition in different cell types. (A) HepG2 or (B) SW480 cancer cells were analyzed upon addition of the indicated components (serum, DMEM, DMEM+DMSO (solvent for ATRA) or all-trans-retinoic acid (ATRA)). Western blot analyses using anti-GPRC5A and anti-Gal-3 antibodies were done on total protein extracts from treated cells. GAPDH was used as a negative control. GPRC5A glycosylated forms are indicated.

### 3.4. GPRC5A endocytosis is mediated by extracellular Gal-3 addition

GPRC5A is an orphan GPCR with no identified ligand and our results show that it interacts with Gal-3. Since Gal-3 mediates endocytic uptake of several glycosylated receptors, we tested whether GPRC5A was internalized following Gal-3 addition. Purified recombinant Gal-3 was added to the cells, and endocytosis assays were performed as previously described [9,14]. In SW480 colorectal cancer cells, GPRC5A expression is increased by retinoic-acid treatment (Figure 3B). Therefore, we treated SW480 cells with or without all-trans-retinoic acid (ATRA) and incubated them in the absence (no Gal-3) or presence (+ Gal-3) of Gal-3 for 25 min at 37°C before assessing GPRC5A localization by immunofluorescence using anti-GPRC5A (supplementary Figure 2). Addition of ATRA did not affect GPRC5A localization, whereas Gal-3 induced its endocytic internalization, evidenced by intracellular puncta observed after 25 min of incubation with Gal-3 (arrows in supplementary Figure 2A, higher magnification in supplementary Figure 2B). These results demonstrate that GPRC5A interacts with Gal-3 and that endocytic internalization is mediated by extracellular addition of Gal-3.

Next, we further analyzed and quantified GPRC5A endocytosis mediated by Gal-3 addition. After 30 minutes of incubation with GRPC5A antibodies on ice, SW480 colorectal cancer cells were incubated at 37°C for 25 minutes in the presence or absence of Gal-3 at a final concentration of 1 µg/ml. Confocal microscopy revealed that treatment with Gal-3 for 25 minutes induced GPRC5A internalization, forming distinct cytosolic puncta (Figure 4A). Internalization of GPRC5A was quantified using ImageJ by measuring fluorescence intensity (Figure 4B), and by counting cells with intracellular fluorescent puncta (Figure 4C). These results indicate that GPRC5A internalization is induced by extracellular Gal-3 addition.

**Figure 4.**
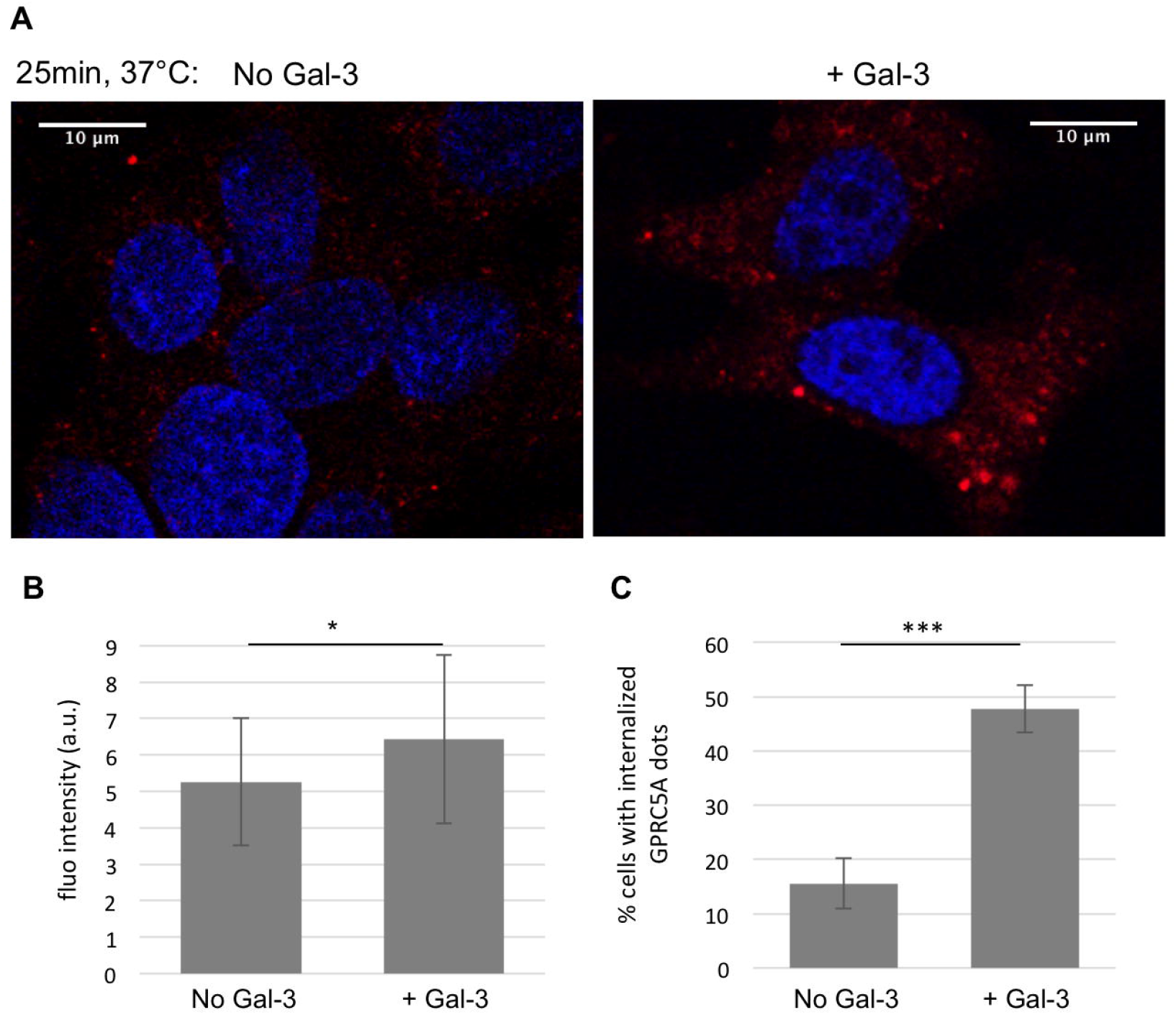
Internalization of GPRC5A upon addition of Gal-3 in SW480 colorectal cancer cells. (A) SW480 colorectal cancer cells were incubated on ice for 30 minutes in presence of anti-GPRC5A (Sigma, HPA046526), then transferred to warm medium containing Gal-3 (1 µg/mL) or without Gal-3 for 25 minutes (+ Gal-3 or No Gal-3). Following washing, fixation, and permeabilization with 0,2% saponin, the GPRC5A antibody was detected using Alexa Fluor-568 conjugated anti rabbit secondary antibody. Slides were mounted using a DAPI-containing mounting medium, and images were acquired a confocal microscope. (B) Red fluorescence intensity was measured using ImageJ for 40 cells per condition. The mean fluorescence intensity is shown in the histogram, and statistical significance was assessed by t-test (^*^p=0,019 < 0,05). (C) The endocytosis experiment described in (A) was performed in triplicate, cells were observed by confocal microscopy, and the percentage of cells with intracellular red fluorescent puncta was quantified. The mean percentage of cells exhibiting GPRC5A endocytosis was calculated with statistical significance assessed by t-test performed (^***^p= 0,0009 <0,001).

## 4. Discussion

GPRC5 receptor family, including GPRC5A, features a short extracellular N-terminal domain and as opposed to other member of GPRC class C, agonists bind to extracellular domain instead of N terminal domain [42,43]. As orphan receptors, GPRC5 members lack identified ligands, but are upregulated by retinoic acid (RA), which has led to their classification as retinoic acid-inducible genes (RAIG) [16]. Herein, we report for the first time that Gal-3, a carbohydrate binding protein, and the orphan G protein coupled receptor GPRC5A physically interact. This interaction was validated in HepG2 hepatocarcinoma and in colorectal SW480 cancer cell lines. Notably, SW480 cells exhibited a higher expression level of both Gal-3 and GPRC5A, which may have a functional significance, as Gal-3 is a key indicator of tumorigenesis, especially in colorectal cancers [44-46]. This finding correlates with the known role of Gal-3 as a regulator of glycosylated proteins, either by their retention in lattices or by internalization [47]. Previous studies have identified various plasma membrane proteins, including CD44 and β1 integrin as Gal-3 binding partners, whose endocytosis is Gal-3 dependent [9,14]. Moreover, recent data show that acute inhibition of Gal-3 strongly decreases α5β1 integrin endocytosis. However, under prolonged Gal-3 inhibition, α5β1 integrin internalization is restored via clathrin-mediated endocytosis [48]. In this study, we observed that GPRC5A internalization was induced by extracellular Gal-3, this cargo being an orphan G-protein coupled receptor whose expression but not its endocytosis is stimulated by treatment with retinoic acid.

Endocytosis pathways enable cells to communicate with their extracellular environment, receive signals and internalize plasma membrane receptors and nutrients. Various endocytic pathways have been described, including clathrin-dependent and clathrin–independent mechanisms, which can operate simultaneously and may provide cellular functional advantages [49]. One such process is the Gal-3 mediated GlycoLipid-Lectin (GL-Lect) endocytosis, which mediates the internalization of glycosylated transmembrane proteins such as CD44, β1 integrin or CD98/SLC3A2 [9,11,14]. GPRC5A is likewise glycosylated and undergoes additional modifications such as palmitoylation [50]. Palmitoylation is a hallmark of GPCRs, with approximately 80% of known GPCRs containing potential cysteine residues for this modification. Experimental evidence confirms palmitoylation for several GPCRs. Palmitoylation affects diverse roles, including G-protein coupling, modulation of endocytosis, and regulation of receptor phosphorylation and desensitization [51]. In transmembrane proteins such as CD44, palmitoylation is essential for lipid raft association, receptor endocytosis, and turnover [52]. Exploring such PTMs could elucidate the mechanisms governing GPRC5A internalization and signaling.

Additionally, interactions between GPRC5B and sphingomyelin synthase 2 (SMS2) have linked sphingolipid metabolism to insulin resistance and obesity [53]. Whether GPRC5A or other GPRC5 subfamily members similarly interact with lipid synthesis pathways remains unknown. Cholesterol, sphingolipids, and glycolipids, vital components of lipid rafts may mediate GPRC5A recruitment and endocytosis [54,55].

Interestingly, exosomes from HT29 colorectal cancer cells contain GPRC5A, Gal-3BP and Tetraspanin 1 (TSPAN1) [56]. In our interactomics data, we identified GPRC5A and Gal-3BP as Gal-3 interacting proteins. Gal-3 is also targeted to exosomes for extracellular release. Indeed, Gal-3 harbors a conserved tetrapeptide motif P(S/T) AP in its N-terminal domain that directly interacts with Tsg101, a component of the endosomal sorting complex required for transport (ESCRT), exosomal secretion of Gal-3 requires Tsg101 and Vps4a [57]. Gal-3 silencing reduces β1-integrin export in exosomes, which is associated with diminished metastatic potential in breast cancer cells [58]. Whether Gal-3 similarly regulates GPRC5A export remains to be determined.

Since GPCRs are among the most successful druggable targets, identifying the endocytosis pathway of the orphan receptor GPRC5A may improve therapeutic and/or prognostic strategies for recurrent and refractory diseases, particularly cancers.

## 5. Conclusions

Our findings reveal GPRC5A as a novel interacting partner of Gal-3. Moreover, we demonstrate that Gal-3 induces GPRC5A internalization. The consequences of this receptor trafficking for cellular functions remain unknown. Comparative analysis with normal cells could clarify whether Gal-3 mediated internalization of GPRC5A is a tumor-specific, and how it influences tumor cell growth, migration potential, or signaling. Finally, the differential expression of glycosylated and non-glycosylated forms of GPRC5A across tumor cell lines highlights the importance of investigating post-translational modifications more closely to unravel their roles in the Gal-3 interaction, internalization mechanisms, and the functional consequences of receptor trafficking. Such studies will improve understanding of the role of the orphan receptor GPRC5A in both normal and cancer cells and may give some hints for potential future therapeutic approaches.

## Author Contributions

“Conceptualization and methodology: SF, AB, SB; Experimental advice: EM, CW, LJ, AB; Performed the experiments: AB, JFM, BR, SB, LB, PH; Performing and Analyzing Mass spectrometry: PH, LK, JC, SZ; Contributed reagents/materials/analysis tools: LJ, CW; Writing: SF, AB, SB; Supervision SF, HR. All authors have read and agreed to the published version of the manuscript.

## Data Availability Statement

The mass spectrometric data were deposited on PRIDE repository with the ProteomeXchange identifier PXD048507.

## Acknowledgments

The authors thank Aline Keilbach (GMGM, CNRS, Université de Strasbourg) for administrative support and Jérémy Thien for help with the ATRA western blot experiments. The colorectal adenocarcinoma cell lines (SW480, Caco-2, DLD1, HT29, HCT116) were a kind gift from Dr Isabelle Gross (CRBS, INSERM, Université de Strasbourg). The ATRA was a kind gift by Norbert B. Ghyselinck (IGBMC, CNRS, Université de Strasbourg).

## Supplementary Figure Legends

**Supplementary Figure 1**. GAL-3 was immunoprecipitated from HepG2 cells protein lysates and samples analyzed by mass spectrometry. Interaction partners were determined by searching against the complete Human proteome set from the SwissProt database. The retained interaction partners were based on the statistical analyses done on the number of spectra (spectral count SC) detected in the negative control immunoprecipitations n=4 omitting GAL-3 antibodies (Ctrl, in green), compared to the GAL-3 immunoprecipitation samples n=4 (IP, in orange). Proteins were considered significantly enriched (significant: Yes) if they met both criteria: log_²_FC ≥ 1 and p-value < 0.05. The proteins having a log²FC ≥ 1 and p-value < 0.15 are also shown, but were not considered as significantly enriched (significant: No). The gene name (GN) and the description are indicated. The full mass spectrometry proteomics data are available in the ProteomeXchange Consortium via the PRIDE partner repository identifier PXD048507 and 10.6019/PXD048507.

**Supplementary Figure 2**. GPRC5A internalization upon Galectin-3 addition but no visible effect of retinoic acid addition. (A) After preincubation with 10-7 M of ATRA or not, SW480 cells were incubated on ice for 30 minutes in presence of anti-GPRC5A before transfer into warm medium containing Galectin-3 (1ug/ml) or not for 25 minutes (No Gal-3 or + Gal-3). After washing, fixation and incubation with 0,2% saponin, the GPRC5A antibody was detected with an AlexaFluor568 coupled secondary antibody. Slides were mounted in DAPI containing mounting medium and observed on a fluorescence microscope, showing internalization upon recombinant Gal-3 addition, but no visible effect of ATRA. (B) SW480 cells treated as in (A), were observed and images shown with a higher magnification.

## References

1. Thiemann, S.; Baum, L.G. Galectins and immune responses-just how do they do those things they do? Annu Rev Immunol 2016, 34, 243–264.

2. Lin, Y.H.; Qiu, D.C.; Chang, W.H.; Yeh, Y.Q.; Jeng, U.S.; Liu, F.T.; Huang, J.R. The intrinsically disordered nterminal domain of galectin-3 dynamically mediates multisite self-association of the protein through fuzzy interactions. The Journal of biological chemistry 2017, 292, 17845–17856.

3. Bouffette, S.; Botez, I.; De Ceuninck, F. Targeting galectin-3 in inflammatory and fibrotic diseases. Trends Pharmacol Sci 2023, 44, 519–531.

4. Boutas, I.; Potiris, A.; Brenner, W.; Lebrecht, A.; Hasenburg, A.; Kalantaridou, S.; Schmidt, M. The expression of galectin-3 in breast cancer and its association with chemoresistance: A systematic review of the literature. Arch Gynecol Obstet 2019, 300, 1113–1120.

5. Jiang, X.N.; Dang, Y.F.; Gong, F.L.; Guo, X.L. Role and regulation mechanism of gal-3 in non-small cell lung cancer and its potential clinical therapeutic significance. Chem Biol Interact 2019, 309, 108724.

6. Portacci, A.; Diaferia, F.; Santomasi, C.; Dragonieri, S.; Boniello, E.; Di Serio, F.; Carpagnano, G.E. Galectin-3 as prognostic biomarker in patients with covid-19 acute respiratory failure. Respir Med 2021, 187, 106556.

7. Li, H.; Li, J.; Xiao, W.; Zhang, Y.; Lv, Y.; Yu, X.; Zheng, J. The therapeutic potential of galectin-3 in the treatment of intrahepatic cholangiocarcinoma patients and those compromised with covid-19. Front Mol Biosci 2021, 8, 666054.

8. Gilson, R.C.; Gunasinghe, S.D.; Johannes, L.; Gaus, K. Galectin-3 modulation of t-cell activation: Mechanisms of membrane remodelling. Prog Lipid Res 2019, 76, 101010.

9. Lakshminarayan, R.; Wunder, C.; Becken, U.; Howes, M.T.; Benzing, C.; Arumugam, S.; Sales, S.; Ariotti, N.; Chambon, V.; Lamaze, C., et al. Galectin-3 drives glycosphingolipid-dependent biogenesis of clathrin-independent carriers. Nature cell biology 2014, 16, 595–606.

10. Ivashenka, A.; Wunder, C.; Chambon, V.; Sandhoff, R.; Jennemann, R.; Dransart, E.; Podsypanina, K.; Lombard, B.; Loew, D.; Lamaze, C., et al. Glycolipid-dependent and lectin-driven transcytosis in mouse enterocytes. Commun Biol 2021, 4, 173.

11. Zhang, C.; Shafaq-Zadah, M.; Pawling, J.; Hesketh, G.G.; Dransart, E.; Pacholczyk, K.; Longo, J.; Gingras, A.C.; Penn, L.Z.; Johannes, L., et al. Slc3a2 n-glycosylation and golgi remodeling regulate slc7a amino acid exchangers and stress mitigation. The Journal of biological chemistry 2023, 299, 105416.

12. MacDonald, E.; Forrester, A.; Valades-Cruz, C.A.; Madsen, T.D.; Hetmanski, J.H.R.; Dransart, E.; Ng, Y.; Godbole, R.; Shp, A.A.; Leconte, L., et al. Growth factor-triggered de-sialylation controls glycolipid-lectin-driven endocytosis. Nature cell biology 2025.

13. Johannes, L.; Shafaq-Zadah, M.; Dransart, E.; Wunder, C.; Leffler, H. Endocytic roles of glycans on proteins and lipids. Cold Spring Harb Perspect Biol 2024, 16.

14. Johannes, L.; Wunder, C.; Shafaq-Zadah, M. Glycolipids and lectins in endocytic uptake processes. J Mol Biol 2016.

15. Hauser, A.S.; Attwood, M.M.; Rask-Andersen, M.; Schioth, H.B.; Gloriam, D.E. Trends in gpcr drug discovery: New agents, targets and indications. Nature reviews. Drug discovery 2017, 16, 829–842.

16. Cheng, Y.; Lotan, R. Molecular cloning and characterization of a novel retinoic acid-inducible gene that encodes a putative g protein-coupled receptor. The Journal of biological chemistry 1998, 273, 35008–35015.

17. Zhao, X.; Stein, K.R.; Chen, V.; Griffin, M.E.; Lairson, L.L.; Hang, H.C. Chemoproteomics reveals microbiota-derived aromatic monoamine agonists for gprc5a. Nat Chem Biol 2023, 19, 1205–1214.

18. Dai, L.; Jin, X.; Liu, Z. Prognostic and clinicopathological significance of gprc5a in various cancers: A systematic review and meta-analysis. PloS one 2021, 16, e0249040.

19. Iglesias González, P.A.; Valdivieso, A.G.; Santa-Coloma, T.A. The g protein-coupled receptor gprc5a-a phorbol ester and retinoic acid-induced orphan receptor with roles in cancer, inflammation, and immunity. Biochemistry and cell biology = Biochimie et biologie cellulaire 2023, 101, 465–480.

20. Debard, S.; Bader, G.; De Craene, J.O.; Enkler, L.; Bar, S.; Laporte, D.; Hammann, P.; Myslinski, E.; Senger, B.; Friant, S., et al. Nonconventional localizations of cytosolic aminoacyl-trna synthetases in yeast and human cells. Methods 2017, 113, 91–104.

21. Ladner, C.L.; Yang, J.; Turner, R.J.; Edwards, R.A. Visible fluorescent detection of proteins in polyacrylamide gels without staining. Analytical biochemistry 2004, 326, 13–20.

22. Waltz, F.; Nguyen, T.T.; Arrive, M.; Bochler, A.; Chicher, J.; Hammann, P.; Kuhn, L.; Quadrado, M.; Mireau, H.; Hashem, Y., et al. Small is big in arabidopsis mitochondrial ribosome. Nat Plants 2019, 5, 106–117.

23. Chicois, C.; Scheer, H.; Garcia, S.; Zuber, H.; Mutterer, J.; Chicher, J.; Hammann, P.; Gagliardi, D.; Garcia, D. The upf1 interactome reveals interaction networks between rna degradation and translation repression factors in arabidopsis. Plant J 2018, 96, 119–132.

24. R-Core-Team. R: A language and environment for statistical computing. https://www.R-project.org/

25. Posit-team. Rstudio: Integrated development environment for r. http://www.posit.co/

26. Wickham H A.M., Bryan J, Chang W, McGowan LD, François R, Grolemund G, Hayes A, Henry L, Hester J, Kuhn M, Pedersen TL, Miller E, Bache SM, Müller K, Ooms J, Robinson D, Seidel DP, Spinu V, Takahashi K, Vaughan D, Wilke C, Woo K, Yutani H. Welcome to the tidyverse. Journal of Open Source Software 2019, 4, 1686.

27. Ginestet, C. Ggplot2: Elegant graphics for data analysis. Journal of the Royal Statistical Society Series A: Statistics in Society 2011, 174, 245–246 %@0964-1998.

28. Wickham, H.; Bryan, J. Readxl: Read excel files. https://readxl.tidyverse.org

29. Schauberger, P.; Walker, A. Openxlsx: Read, write and edit xlsx files. https://ycphs.github.io/openxlsx/index.html

30. Slowikowski, K. Ggrepel: Automatically position non-overlapping text labels with ‘ggplot2’. https://ggrepel.slowkow.com/

31. Perez-Riverol, Y.; Bai, J.; Bandla, C.; García-Seisdedos, D.; Hewapathirana, S.; Kamatchinathan, S.; Kundu, D.J.; Prakash, A.; Frericks-Zipper, A.; Eisenacher, M., et al. The pride database resources in 2022: A hub for mass spectrometry-based proteomics evidences. Nucleic Acids Res 2022, 50, D543–d552.

32. Stoetzel, C.; Bar, S.; De Craene, J.O.; Scheidecker, S.; Etard, C.; Chicher, J.; Reck, J.R.; Perrault, I.; Geoffroy, V.; Chennen, K., et al. A mutation in vps15 (pik3r4) causes a ciliopathy and affects ift20 release from the cis-golgi. Nature communications 2016, 7, 13586.

33. Lin, T.W.; Chang, H.T.; Chen, C.H.; Chen, C.H.; Lin, S.W.; Hsu, T.L.; Wong, C.H. Galectin-3 binding protein and galectin-1 interaction in breast cancer cell aggregation and metastasis. J Am Chem Soc 2015, 137, 9685–9693.

34. Mathew, M.P.; Donaldson, J.G. Distinct cargo-specific response landscapes underpin the complex and nuanced role of galectin-glycan interactions in clathrin-independent endocytosis. The Journal of biological chemistry 2018, 293, 7222–7237.

35. Joeh, E.; O’Leary, T.; Li, W.; Hawkins, R.; Hung, J.R.; Parker, C.G.; Huang, M.L. Mapping glycan-mediated galectin-3 interactions by live cell proximity labeling. Proceedings of the National Academy of Sciences of the United States of America 2020, 117, 27329–27338.

36. Couves, E.C.; Gardner, S.; Voisin, T.B.; Bickel, J.K.; Stansfeld, P.J.; Tate, E.W.; Bubeck, D. Structural basis for membrane attack complex inhibition by cd59. Nature communications 2023, 14, 890.

37. Xiong, L.; Edwards, C.K., 3rd; Zhou, L. The biological function and clinical utilization of cd147 in human diseases: A review of the current scientific literature. International journal of molecular sciences 2014, 15, 17411–17441.

38. Capone, E.; Iacobelli, S.; Sala, G. Role of galectin 3 binding protein in cancer progression: A potential novel therapeutic target. Journal of translational medicine 2021, 19, 405.

39. Priglinger, C.S.; Szober, C.M.; Priglinger, S.G.; Merl, J.; Euler, K.N.; Kernt, M.; Gondi, G.; Behler, J.; Geerlof, A.; Kampik, A., et al. Galectin-3 induces clustering of cd147 and integrin-β1 transmembrane glycoprotein receptors on the rpe cell surface. PloS one 2013, 8, e70011.

40. Chen, Y.; Deng, J.; Fujimoto, J.; Kadara, H.; Men, T.; Lotan, D.; Lotan, R. Gprc5a deletion enhances the transformed phenotype in normal and malignant lung epithelial cells by eliciting persistent stat3 signaling induced by autocrine leukemia inhibitory factor. Cancer Res 2010, 70, 8917–8926.

41. Greenhough, A.; Bagley, C.; Heesom, K.J.; Gurevich, D.B.; Gay, D.; Bond, M.; Collard, T.J.; Paraskeva, C.; Martin, P.; Sansom, O.J., et al. Cancer cell adaptation to hypoxia involves a hif-gprc5a-yap axis. EMBO Mol Med 2018, 10.

42. Bräuner-Osborne, H.; Jensen, A.A.; Sheppard, P.O.; Brodin, B.; Krogsgaard-Larsen, P.; O’Hara, P. Cloning and characterization of a human orphan family c g-protein coupled receptor gprc5d. Biochimica et biophysica acta 2001, 1518, 237–248.

43. Lagerström, M.C.; Schiöth, H.B. Structural diversity of g protein-coupled receptors and significance for drug discovery. Nature reviews. Drug discovery 2008, 7, 339–357.

44. Nangia-Makker, P.; Balan, V.; Raz, A. Regulation of tumor progression by extracellular galectin-3. Cancer Microenviron 2008, 1, 43–51.

45. Wang, Y.; Liu, S.; Tian, Y.; Wang, Y.; Zhang, Q.; Zhou, X.; Meng, X.; Song, N. Prognostic role of galectin-3 expression in patients with solid tumors: A meta-analysis of 36 eligible studies. Cancer cell international 2018, 18, 172.

46. Aureli, A.; Del Corno, M.; Marziani, B.; Gessani, S.; Conti, L. Highlights on the role of galectin-3 in colorectal cancer and the preventive/therapeutic potential of food-derived inhibitors. Cancers (Basel) 2022, 15. 47.

47. Mukherjee, M.M.; Biesbrock, D.; Hanover, J.A. Galectin-3: Integrator of signaling via hexosamine flux. Biomolecules 2025, 15, 1028.

48. Shafaq-Zadah, M.; Dransart, E.; Mani, S.K.; Sampaio, J.L.; Bouidghaghen, L.; Nilsson, U.J.; Leffler, H.; Johannes, L. Exploration into galectin-3 driven endocytosis and lattices. Biomolecules 2024, 14.

49. Thottacherry, J.J.; Sathe, M.; Prabhakara, C.; Mayor, S. Spoiled for choice: Diverse endocytic pathways function at the cell surface. Annual review of cell and developmental biology 2019, 35, 55–84.

50. Yang, W.; Di Vizio, D.; Kirchner, M.; Steen, H.; Freeman, M.R. Proteome scale characterization of human s-acylated proteins in lipid raft-enriched and non-raft membranes. Molecular & cellular proteomics : MCP 2010, 9, 54–70.

51. Chalhoub, G.; McCormick, P.J. Palmitoylation and g-protein coupled receptors. Prog Mol Biol Transl Sci 2022, 193, 195–211.

52. Thankamony, S.P.; Knudson, W. Acylation of cd44 and its association with lipid rafts are required for receptor and hyaluronan endocytosis. The Journal of biological chemistry 2006, 281, 34601–34609.

53. Kim, Y.J.; Greimel, P.; Hirabayashi, Y. Gprc5b-mediated sphingomyelin synthase 2 phosphorylation plays a critical role in insulin resistance. iScience 2018, 8, 250–266.

54. Rog, T.; Vattulainen, I. Cholesterol, sphingolipids, and glycolipids: What do we know about their role in raft-like membranes? Chem Phys Lipids 2014, 184, 82–104.

55. Hirabayashi, Y.; Kim, Y.J. Roles of gprc5 family proteins: Focusing on gprc5b and lipid-mediated signalling. J Biochem 2020, 167, 541–547.

56. Lee, C.H.; Im, E.J.; Moon, P.G.; Baek, M.C. Discovery of a diagnostic biomarker for colon cancer through proteomic profiling of small extracellular vesicles. BMC Cancer 2018, 18, 1058.

57. Bänfer, S.; Jacob, R. Galectins in intra- and extracellular vesicles. Biomolecules 2020, 10.

58. Zhang, D.X.; Dang, X.T.T.; Vu, L.T.; Lim, C.M.H.; Yeo, E.Y.M.; Lam, B.W.S.; Leong, S.M.; Omar, N.; Putti, T.C.; Yeh, Y.C., et al. Avβ1 integrin is enriched in extracellular vesicles of metastatic breast cancer cells: A mechanism mediated by galectin-3. Journal of extracellular vesicles 2022, 11, e12234.

