## Supplementary figures and images for "Galectin-3 Mediated Endocytosis of the Orphan G-Protein-Coupled Receptor GPRC5A"

### Supplementary Figure 1

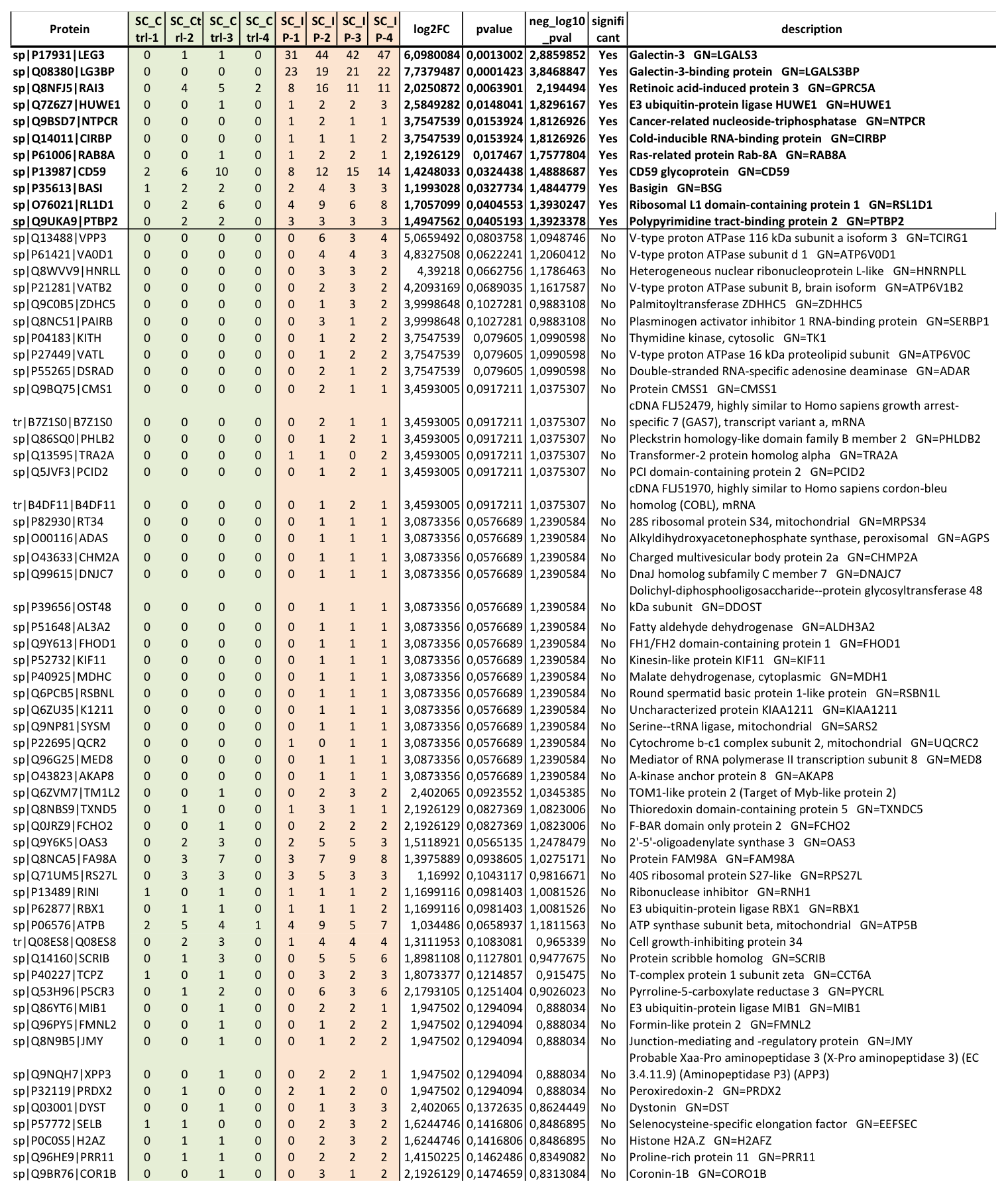

### Supplementary Figure 2

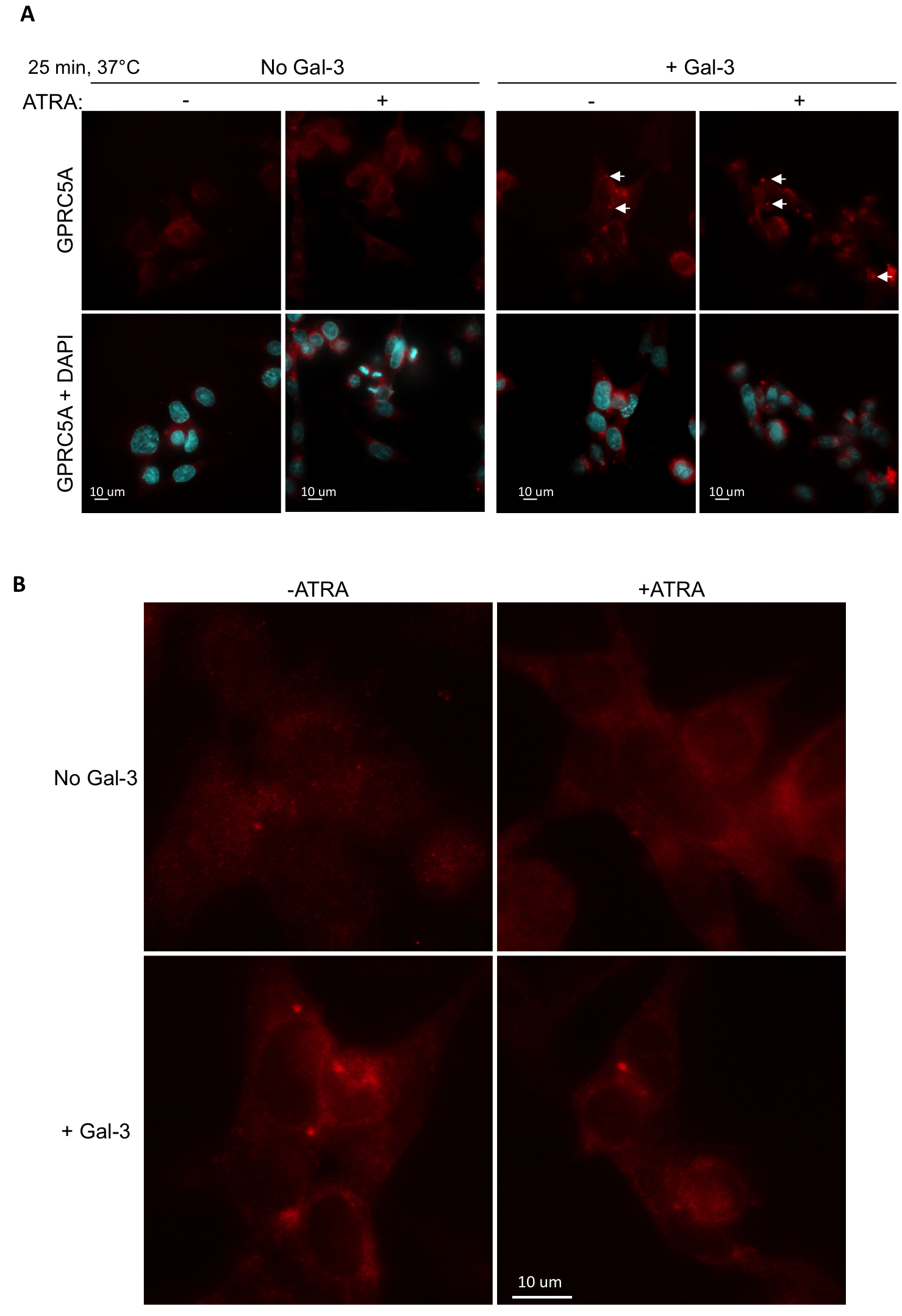
